# Structural and computational insights into the SARS-CoV-2 Omicron RBD-ACE2 interaction

**DOI:** 10.1101/2022.01.03.474855

**Authors:** Jun Lan, Xinheng He, Yifei Ren, Ziyi Wang, Huan Zhou, Shilong Fan, Chenyou Zhu, Dongsheng Liu, Bin Shao, Tie-Yan Liu, Qisheng Wang, Linqi Zhang, Jiwan Ge, Tong Wang, Xinquan Wang

**Author notes:** These authors contributed equally to this work (Jun Lan, Xinheng He, Yifei Ren, Ziyi Wang). Correspondence, (L.Z.), (J. G.), (T.W.), (X.W.).

## Abstract

Since SARS-CoV-2 Omicron variant (B.1.1.529) was reported in November 2021, it has quickly spread to many countries and outcompeted the globally dominant Delta variant in several countries. The Omicron variant contains the largest number of mutations to date, with 32 mutations located at spike (S) glycoprotein, which raised great concern for its enhanced viral fitness and immune escape^[1–4]^. In this study, we reported the crystal structure of the receptor binding domain (RBD) of Omicron variant S glycoprotein bound to human ACE2 at a resolution of 2.6 Å. Structural comparison, molecular dynamics simulation and binding free energy calculation collectively identified four key mutations (S477N, G496S, Q498R and N501Y) for the enhanced binding of ACE2 by the Omicron RBD compared to the WT RBD. Representative states of the WT and Omicron RBD-ACE2 systems were identified by Markov State Model, which provides a dynamic explanation for the enhanced binding of Omicron RBD. The effects of the mutations in the RBD for antibody recognition were analyzed, especially for the S371L/S373P/S375F substitutions significantly changing the local conformation of the residing loop to deactivate several class IV neutralizing antibodies.

---

Since the first documented cases of the SARS-CoV-2 infection in Wuhan, China in late 2019, the COVID-19 pandemic has been posing a severe threat to the global public health, with more than 286 million infections and over 5.4 million deaths around the world (https://www.who.int/emergencies/diseases/novel-coronavirus-2019/situation-reports/). The global fight against the COVID-19 is still in great uncertainty due to the emergence of SARS-CoV-2 variants, especially the variants of concern (VOC) with changed pathogenicity, increased transmissibility and resistance to convalescent/vaccination sera and monoclonal neutralizing antibodies^[5–11]^. In 2020, the first VOC Alpha (B.1.1.7) was identified in the United Kingdom, followed by the Beta (B.1.351) in South Africa and Gamma (P.1) in Brazil. These three VOCs mainly circulated in their identified and neighboring countries. In contrast, the Delta variant first detected in India in late 2020 quickly spread to nearly all countries and has become the global dominant VOC in 2021. In November 2021, the newest Omicron variant (B.1.1.529) was reported from South Africa, and the World Health Organization (WHO) immediately designated it as the fifth VOC due to its close to 40 mutations in the spike (S) glycoprotein, at least three times more than the number found in previous four VOCs^[12–15]^. Notably, the Omicron variant has unprecedented 15 mutations in the RBD that recognizes host ACE2 receptor and is also the major target of potent neutralizing antibodies, whereas the Alpha, Beta, Gamma and Delta variants have only 1, 3, 3 and 2 mutations in the RBD, respectively. So many mutations occurred simultaneously in the RBD would have a significant impact on the infectivity and transmissibility of the Omicron variant and the protective effects of current approved vaccines and therapeutic antibodies^[1–3]^. A detailed description of the Omicron RBD and its receptor and antibody interactions is essential for fully understanding the infection and immune escape of the Omicron variant at the molecular level.

The Omicron RBD (residues Arg319-Phe541) and the N-terminal peptidase domain of ACE2 (residues Ser19-Asp615) were expressed in Hi5 insect cells and purified by Ni-NTA affinity and gel-filtration. The complex structure was determined by molecular replacement and refined at 2.6 Å resolution to final *R*_work_ and *R*_free_ factors of 19.2% and 23.1%, respectively (**Extended Data Table 1**). The final model contains residues from Thr333 to Gly526 of the Omicron RBD and residues from Pro18 to Asp615 of the ACE2 N-terminal peptidase domain. The overall structure of Omicron RBD-ACE2 complex is highly similar to that of the WT RBD-ACE2 complex^[16]^, with an r.m.s.d. of 0.209 Å for 730 aligned Cα atoms (**Fig. 1a**). There are 15 mutations in the Omicron RBD (G339D, S371L, S373P, S375F, K417N, N440K, G446S, S477N, T478K, E484A, Q493K/R, G496S, Q498R, N501Y, Y505H), with ten mutations in the RBM for ACE2 binding and five mutations in the core subdomain (**Fig. 1a**). The Omicron RBD in our study contains the Q493K substitution, which was reported in the beginning and was recently replaced by Q493R mutation^[17]^. Three mutations S371L, S373P and S375F in the core subdomain are clustered at a hairpin loop (residues Y369-C379), resulting in a main-chain conformational change compared to the WT and other mutated RBD structures (**Fig. 1b**).

**Figure 1.**
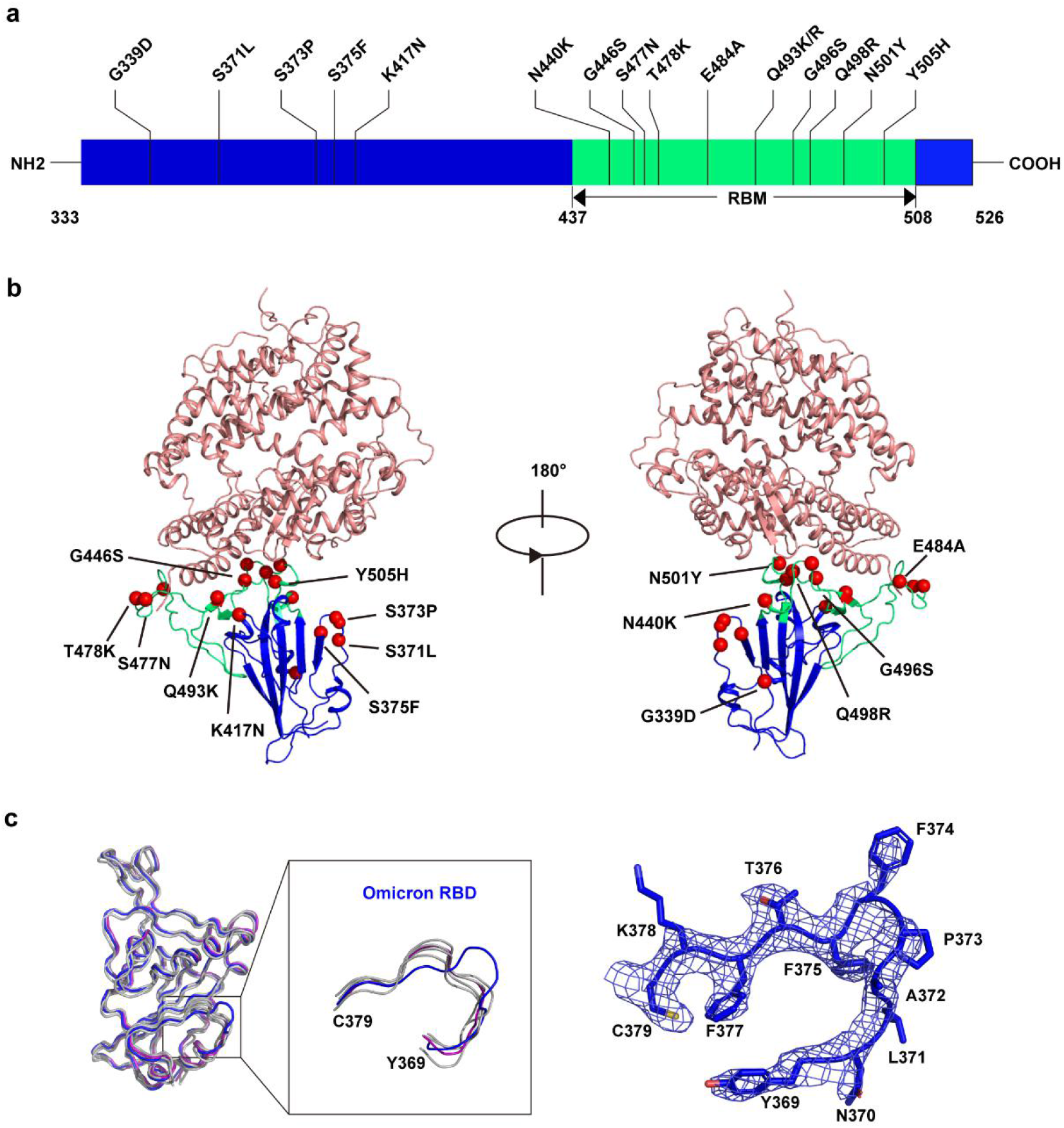
Overall structure of the Omicron RBD bound to ACE2. (**a**) Upper: substitution of amino acid residues on the Omicron RBD. Ten residues are located at the RBM region (green) and five residues are located at the core subdomain (blue). Lower: complex structure of the Omicron RBD and ACE2. RBD core subdomain is shown in blue, and the RBM is shown in green. ACE2 is shown in salmon. Mutation residues are colored in red balls. (**b**) Alignment of the Omicron RBD structure with other WT and mutated RBDs structures previously reported with resolutions higher than 3.2 Å (PDB ID 6M0J, 7E7Y, 7NX6, 7NXC and 7R6W). The Omicron RBD is colored in blue. The WT RBD is colored in purple. The other RBDs are shown in gray. 2*F*o-*F*c electron density map of the Omicron RBD hairpin loop (Y369-C379) contoured at 1.5σ is shown.

The ten mutations in the RBM do not significantly change the overall binding mode of the Omicron RBD with ACE2, compared to that between WT RBD and ACE2. With a distance cut-off of 4 Å, the numbers of contacting residues at the interface are nearly the same (16 Omicron RBD residues with 21 ACE2 residues; 17 WT RBD residues with 20 ACE2 residues) (**Extended Data Table 2**). The Omicron RBD-ACE2 complex also has a slightly increased buried surface at the interface (~1,748 Å^2^), compared to the buried surface of 1,687 Å^2^ in the WT RBD-ACE2 complex. Although the overall binding mode is highly conserved, a slight difference in the binding affinity was observed between the Omicron and WT RBDs to ACE2. Using the surface plasmon resonance (SPR) method, the measured binding affinity between the Omicron RBD and ACE2 is approximately 2.8 nM, exhibiting a 2-fold increased binding compared to the WT RBD (K_D_= 5.9 nM) (**Fig. 2a**).

**Figure 2.**
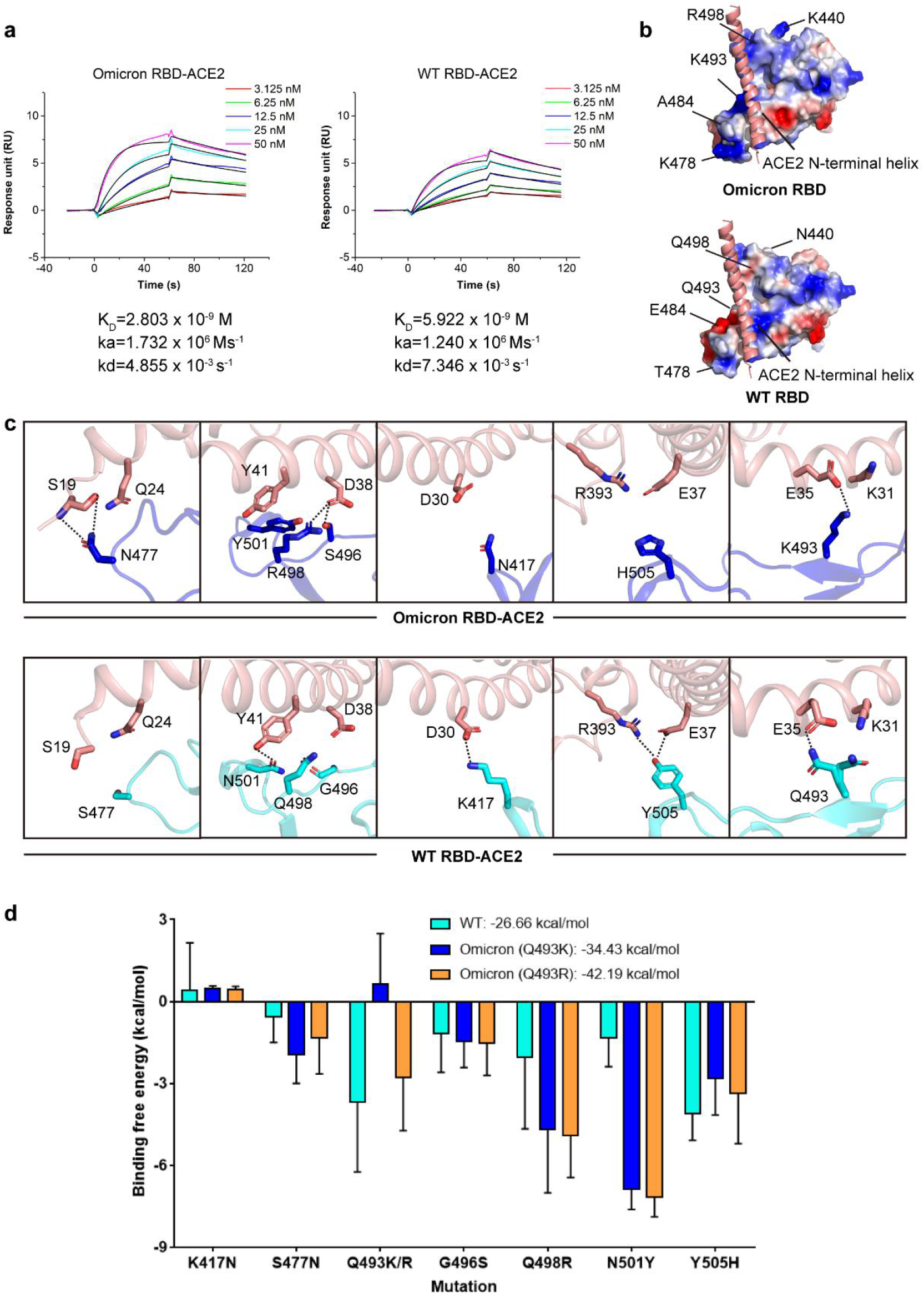
Comparison of the binding to ACE2 between Omicron and WT RBDs. (**a**) SPR curves of ACE2 with Omicron RBD (left) and WT RBD (right). Experimental curves are shown as colored lines and the fitting curves are colored in black. (**b**) Electrostatic potential maps of the Omicron RBD (upper) and WT RBD (lower). The N-terminal helix is shown in a salmon ribbon. The positions of mutation residues are indicated by a black line. (**c**) Change of interactions with ACE2 between Omicron RBD (upper panel) and WT RBD (down panel) at the S477N, G496S, Q498R, N501Y, K417N, Y505H and Q493 sites. Omicron RBD, WT RBD and ACE2 are shown in blue, cyan and salmon, respectively. Contacting residues are shown as sticks. Hydrogen bonds are represented by dashed lines. (**d**) The energy decomposition of key interface residues for Omicron and WT RBD. WT, Omicron (Q493K), and Omicron (Q493R) are colored in cyan, blue, and orange, respectively.

To reveal the molecular details in determining the change of binding affinity, we compared and analyzed the variations of interactions between the Omicron and WT RBDs in their binding to the ACE2 receptor. The first variation is about the surface electrostatic potential of the Omicron RBD. Compared to the WT RBD, Omicron RBD has a more positive electrostatic potential in the surface for ACE2 binding due to the substitutions to basic residues including N440K, T478K, Q493K and Q498R and the loss of acidic residue E484A (**Fig. 2b**). In addition, the Omicron RBD obtains newly formed interactions at several sites. One is at the RBD 477 site, in which the WT RBD S477 is not in contact with ACE2, whereas the N477 in the Omicron RBD forms interactions with ACE2 S19 and Q24 (**Fig. 2c**). A more significant one is around a cluster of RBD 496, 498 and 501 sites. At the WT RBD-ACE2 interface, van del Waals interactions are only observed from WT RBD Q498 and N501 to ACE2 Y41 and Q42. Due to G496S, Q498R and N501Y mutations in the Omicron RBD, newly formed interactions occur between Omicron RBD residues (S496, R498 and Y501) and ACE2 residues (D38, Y41 and Q42). Specifically, S496 forms a hydrogen bond with ACE2 D38; R498 forms a new hydrogen bond and salt bridge with ACE2 D38; and additional π-π interaction is formed between Omicron RBD Y501 and ACE2 Y41 (**Fig. 2c, Extended Data Table 2 and Extended Data Table 3**). In the meantime, the mutations also result in the loss of interactions between the Omicron RBD and ACE2 in comparison to the WT RBD. No interaction is observed between Omicron RBD N417 and ACE2 D30, whereas a salt bridge is formed between WT RBD K417 and ACE2 D30. The Y505H mutation in Omicron RBD disrupts previous hydrogen-bonding interactions between WT RBD Y505 and ACE2 E37 and R393 (**Fig. 2c, Extended Data Table 2 and Extended Data Table 3**). Regarding the RBD 493 site, the Q493K/R substitution results in a newly formed salt bridge with ACE2 E35, while also disrupting previous interactions of Q49 with ACE2 K31 (**Fig. 2c, Extended Data Table 2 and Extended Data Table 3**). We also utilized Molecular Dynamics (MD) simulation and Molecular Mechanics Generalized Born Surface Area (MM/GBSA) method to estimate the binding free energies (**Fig. 2d)**. The binding free energies of the Omicron and WT RBDs to ACE2 are −42.19 ± 6.61 and −26.66 ± 9.78 kcal/mol, respectively, which also supports the enhanced binding of the Omicron RBD to ACE2. We also analyzed the contribution of each mutation in the binding free energy change (**Fig. 2d)**. Four mutations including S477N, G496S, Q498R and N501Y have positive impacts on the binding of the Omicron RBD to ACE2 by further reducing the free energy by 0.77, 0.36, 2.85 and 5.84 kcal/mol respectively, which is consistent with the above described newly formed interactions of at these four sites, especially around the 496, 498 and 501 cluster. We also observed the negative impacts of Q493K/R and Y505H mutations. The negative impact of the mutation at 493 site is a little unexpected because a new salt bridge to ACE2 E35 is formed by Q493K/R mutation. The loss of Q493 to ACE2 K31 interaction and the close position of positively charged ACE2 K31 and Omicron RBD K493 or R493 may exceed the positive effect of the salt bridge to ACE2 E35 and finally determine the net negative impact of Q493K/R mutation.

To explore the variations between the WT and Omicron RBD-ACE2 interactions from a dynamic view, we run microsecond level of MD simulations for the WT and Omicron systems and utilized Markov State Model (MSM) to identify representative states and illustrate the transition process among them. Time-lagged independent component (tIC) analysis by projecting the conformations onto the low-dimensional surface showed that the WT and Omicron RBD-ACE2 interface contacts have different distributions in the first two components, tIC 1 and tIC 2 (**Fig. 3a)**. With a novel lag time chosen algorithm recently designed by us, all conformations were first clustered into microstates and then into macrostates, which are representative conformations for a set of points on the energy landscape (**Fig. 3a**). Finally, three macrostates 1, 2, and 3 were sorted out according to their proportions in all conformations (**Fig. 3a**). The state 3 of the WT and Omicron locate at a similar position around (0, −1) on the energy landscape. Although the positions of state 2 in WT and Omicron are also similar, the WT system shows a larger value (> 2.0) on the tIC 2 dimension. The state 1 of WT locates at (−2, −1) of the energy landscape that is not visited by Omicron, and the coordinate (2,1) of the state 1 in the Omicron is also not touched by the WT either. We further extracted the representative structures for each macrostate and analyzed the transition process between each pair of macrostates by transition path theory (**Fig. 3b and Extended Data Fig. 1-2**). In the WT system, state 3 is the dominant one and the transition from states 1 and 2 to it is very quick (< 1 ns). State 2 is the second large macrostate which covers around 8.7 % conformational space and is relatively easy to be reached from states 1 and 3 (both are 4.90 ns). In the Omicron system, the distributions of states 1, 2 and 3 are relatively balanced with a proportion of 16.6%, 31.1%, and 52.3%, respectively, and similar to the WT system, state 3 can be quickly reached from states 1 and 2 with 1.07 ns and 0.89 ns, respectively. Furthermore, it can also easily transfer back to the other two states (4.12 ns to state 1 and 1.70 ns to state 2).

**Figure 3.**
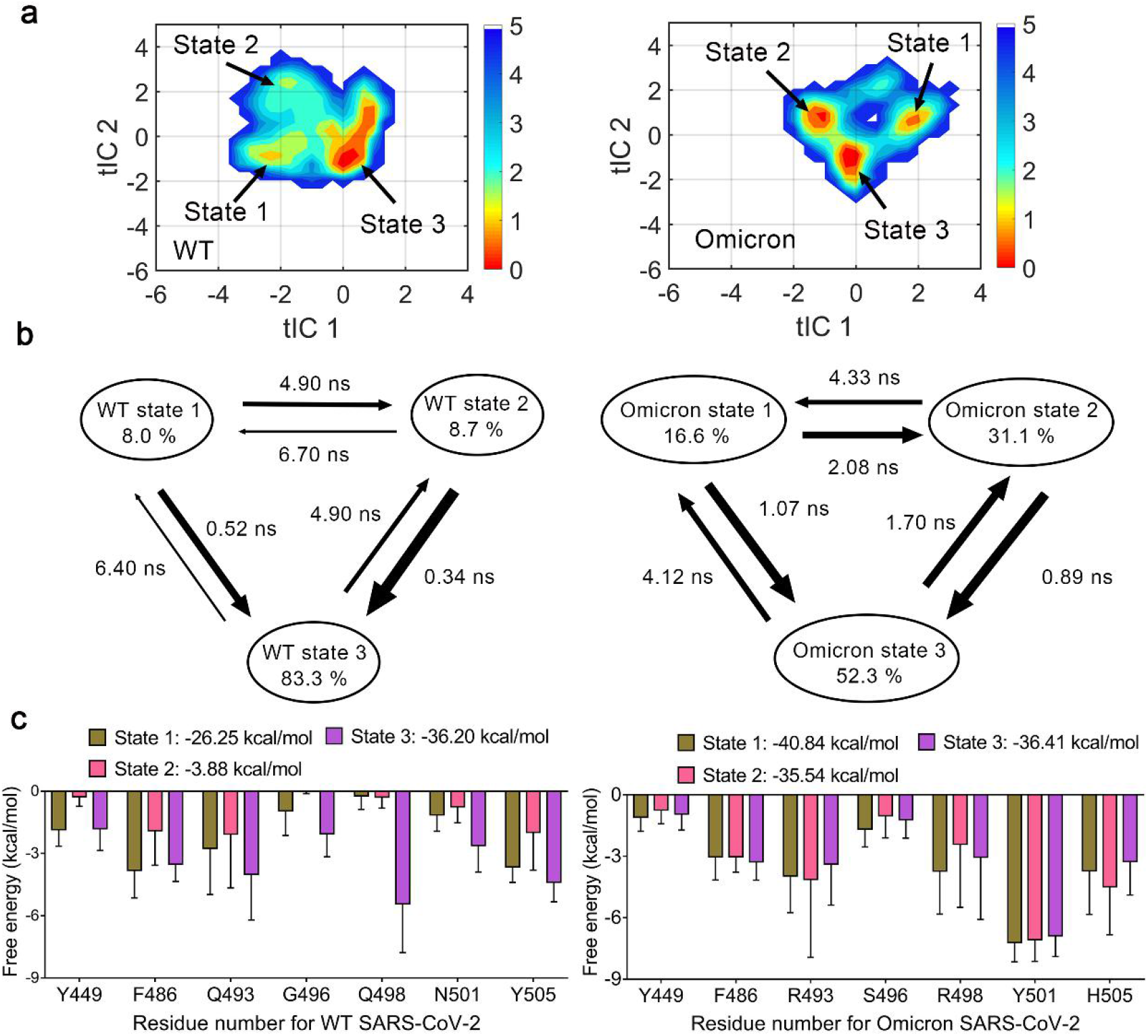
Representative states and key residues identified and analyzed at the interacting interface between the RBD and ACE2. (**a**) The free energy landscape of WT and Omicron system. The tIC 1 and tIC 2 were constructed according to the contacts between residue pairs of RBD and ACE2. The corresponding macrostates are labeled by arrows. (**b**) The proportions for each macrostate and transition time between states in WT and Omicron systems. The thickness of arrows is proportional to the reciprocal of transition time. (**c**) The energy decomposition of key residues in each macrostate. Residues in state1, 2 and 3 are colored in olive, pink and purple, respectively.

Considering the differences in the distributions and kinetics of representative states between the WT and omicron systems, we further evaluated the binding abilities for each macrostate by employing MM/GBSA (**Fig. 3c**). For the WT simulation system, state 2 is the least suitable conformation for ACE2 binding in all macrostates (ΔG = −3.88 ± 19.64 kcal/mol), state 1 has a better binding ability (ΔG = −26.25 ± 9.76 kcal/mol), and the dominant state 3 is the best-fit conformation for ACE2 binding (Δ G = −36.20 ± 8.64 kcal/mol). As for the Omicron system, state 1 shows the highest binding affinity with ACE2 (ΔG = −40.84 ± 6.93 kcal/mol), and states 2 and 3 have similar binding affinities (ΔG =-35.54 ± 8.11 and-36.41 ± 10.26 kcal/mol, respectively). Notably, in the WT system, the dominant state 3 has the lowest binding free energy, whereas the other two minor states 1 and 2 have significantly higher binding free energies. In contrast, all three macrostates in the Omicron system have low binding free energies and the differences among them are not as significant as those in the WT system. We also decomposed and compared the binding free energy to residues and picked up the residues with highly different energy values among macrostates in WT system. As shown in **Fig. 3c and Extended Data Fig. 3-5**, Y449, F486, Q493, G496, Q498, N501 and Y505 decrease the contributions of the binding free energy in WT state 1 but the mutated residues in the Omicron RBD have more favorable ACE2 interactions at the interface and thus rescue the binding free energy. Taken together, the dynamic transition process in the RBD-ACE2 interface suggests that the WT RBD samples conformations disfavoring ACE2 binding, whereas the mutations including Q493R, G496S, Q498R, N501Y and Y505H prevent the Omicron RBD to sample the disfavoring conformations, thereby enhancing its binding to ACE2.

In addition to the impact on ACE2 binding, the mutations in the RBD enable the Omicron variant to escape from antibody recognition and neutralization. Several recent studies have evaluated the neutralizing activities of antibodies against the Omicron variant^[1,2,17,18]^. Their results showed that most of the studied antibodies, especially Class I and Class II antibodies, lost their neutralizing activities due to mutations in the epitopes such as K417N and E484A, which is consistent with previous studies^[19–21]^. Most of the studied Class III and Class IV antibodies still maintained neutralizing activities. However, a subset of Class III antibodies were reported to be sensitive to the G339D, N440K, and G446S mutations in the Omicron RBD^[1,2]^. We paid special attention to the S371L/S373P/S375F mutations within the epitopes of a subset of Class IV antibodies including S2X35, S304, SA24, H104 and CR3022^[22–25]^ (**Fig. 4a-c**). Indeed, the significantly reduced neutralization activities have been reported for S2X35 and S304 against the Omicron variant^[18]^. Structural studies have revealed hydrogen-bonding interactions occurred between these two antibodies and the hairpin loop (residue Y369-C379) (**Fig. 4d and 4e**). 16 hydrogen bonds are formed at the interface of RBD and these two antibodies. Specially, half of the hydrogen bonds are formed with main chain atoms of RBD residues and the remaining ones are formed between the side chain atoms (**Fig. 4d, 4e**). The S371L/S373P/S375F substitutions not only changed the side chains of residues, but also induced a main-chain conformational change, which would disrupt the specific binding of the antibodies to the hairpin loop (residue Y369-C379). It is expected that antibodies including the hairpin loop in their recognizing epitopes would also be impaired by the Omicron RBD S371L/S373P/S375F substitutions for their binding and neutralization.

**Figure 4.**
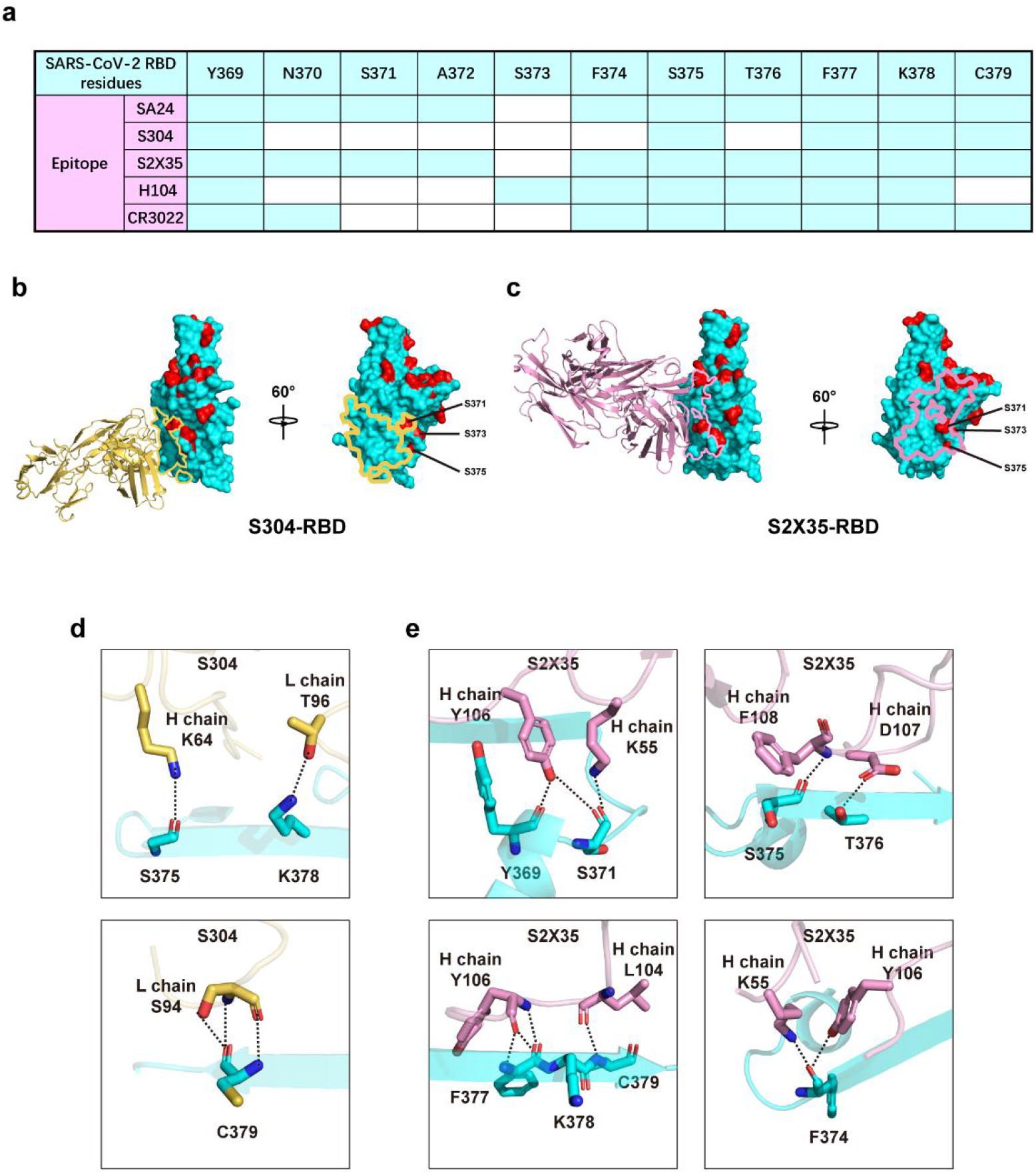
Structural basis of S371L/S373P/S375F mutations for escaping the neutralization of Class IV antibodies. (**a**) The binding residues of Class IV antibodies (SA24, S304, S2X35, H104 and CR3022) at the hairpin loop. The residues recognized by antibodies is labeled with cyan box. (**b, c**) The binding mode and epitopes of representative Class IV antibodies on RBD; S304 Fab-RBD complex (**b**) and S2X35 Fab-RBD complex (**c**) are shown, with RBD shown as cyan surface. S304 Fab and S2X35 Fab is shown as cartoon with yellow and pink, respectively. The epitope of S304 and S2X35 on RBD is shown with yellow and pink lines, respectively. The mutations of Omicron variant are shown with red surface. (**d, e**) Interactions between the S304 (**d**) or S2X35 (**e**) and SARS-CoV-2 RBD. The hydrogen bonds are displayed with black dashed lines.

The mutations on the S glycoprotein of various SARS-CoV-2 variants determine the viral fitness by mainly affecting cell infection and immune escape. For enhancing the fitness during virus-host evolution, it is expected that mutations of the RBD residues, especially residues in the RBM for both ACE2 binding and antibody recognition need to be balanced to keep or increase ACE2 binding while to disrupt antibody recognition at the same time. Previous SARS-CoV-2 VOCs including Alpha, Beta, Gamma and Delta only have 1-3 mutated residues in the RBD, whereas the newest Omicron variant has a surprising 15 mutations in the RBD. Therefore, understanding the effects of these mutated RBD residues on ACE2 binding and antibody recognition at an atomic level are very important. In this study, we utilized crystal structure determination and computational analysis to analyze the Omicron RBD-ACE2 interaction from both static and dynamic viewpoints. Our results elucidated the important roles of new mutations S477N, G496S and Q498R for the enhanced binding of the Omicron RBD to ACE2. Our dynamic analysis also revealed that these mutations on the Omicron RBD could collectively affect the conformational sampling of the RBD for ACE2 binding by avoiding disfavoring conformations as seen in the WT RBD. Previous studies have provided amounts of information about the epitopes of potent neutralizing antibodies. Through structural comparisons of the different complex structures of RBD and neutralizing antibodies, we were also able to reveal structural basis for the immune escape of the Omicron variant, especially for the clustered S371L/S373P/S375F substitutions that deactivate several Class IV neutralizing antibodies by inducing changes of both main-chain and side-chain conformations.

## Methods

### Protein expression and purification

The SARS-CoV-2 Omicron RBD and the N-terminal peptidase domain of ACE2 were expressed using the Bac-to-Bac baculovirus system (Invitrogen) as previously stated. Briefly, The SARS-CoV-2 Omicron RBD (residues Thr333–Gly526) with an N-terminal gp67 signal peptide for secretion and a C-terminal 6×His tag for purification was expressed using Hi5 cells and purified by Ni-NTA resin and gel filtration chromatography (GE Healthcare) in HBS buffer (10 mM HEPES, pH 7.2, 150 mM NaCl).

The N-terminal peptidase domain of ACE2 (residues Ser19–Asp615) was expressed and purified by essentially the same protocol as used for the SARS-CoV-2 Omicron RBD. To obtain the SARS-CoV-2 Omicron RBD and human ACE2 complex, ACE2 was incubated with the SARS-CoV-2 Omicron RBD for 1 h on ice in HBS buffer, and the mixture was then subjected to gel filtration chromatography. Fractions containing the complex were pooled and concentrated to 13 mg ml^-1^.

### Crystallization and data collection

Crystals were successfully grown at room temperature in sitting drops, over wells containing 0.2 M L-Proline, 0.1 M HEPES pH 7.5, 10% w/v Polyethylene glycol 3,350. Crystals were collected, soaked briefly in 0.2 M L-Proline, 0.1 M HEPES pH 7.5, 10% w/v Polyethylene glycol 3,350 and 20% glycerol, and were subsequently flash-frozen in liquid nitrogen. Diffraction data were collected at 100 K and a wavelength of 1.07180 Å at the BL02U1 beam line of the Shanghai Synchrotron Research Facility. Diffraction data were processed using the HKL3000 software^[26]^ and the data-processing statistics are listed in Extended Data Table 1.

### Structure determination and refinement

The structure was determined using the molecular replacement method with PHASER in the CCP4 suite^[27]^. The search models used included the ACE2 extracellular domain and SARS-CoV-2 RBD (PDB: 6M0J). Subsequent model building and refinement were performed using COOT^[28]^ and PHENIX^[29]^, respectively. Final Ramachandran statistics: 96.95% favored, 3.05% allowed and 0.00% outliers for the final structure. The structure refinement statistics are listed in Extended Data Table 1. All structure figures were generated with PyMol^[30]^.

### Surface plasmon resonance

Binding kinetics of ACE2 and SARS-CoV-2 RBDs were determined by surface plasmon resonance using a Biacore S200 (GE Healthcare). All experiments were performed in a running buffer composed of 10 mM HEPES, pH 7.2, 150 mM NaCl, and 0.005% Tween-20 (v/v). ACE2 was immobilized on a CM5 sensor chip (GE Healthcare) to a level of ~500 response units. A 2-fold dilution series ranging from 50 to 3.125 nM of the SARS-CoV-2 WT RBD and omicron RBD were injected at a flow rate of 30 μl/min (association 60s, dissociation 180s), and the immobilized ACE2 was regenerated using 5mM NaOH for 10s. The resulting data were fit to a 1:1 binding model using Biacore Evaluation Software (GE Healthcare).

### MD simulations

The Omicron RBD-ACE2 structure and WT RBD-ACE2 complex (PDB ID: 6M0J) were used to build the MD simulation systems. Besides, K493 in Omicron RBD-ACE2 structure was mutated to R493 to follow the recent sequence of Omicron. Therefore, three simulation systems including WT, Omicron (Q493K) and Omicron(Q493R) were constructed. To keep the consistency among simulation systems, S19-D615 in ACE2 and T333-G526 in RBD were maintained in WT and the two Omicron systems.

The FF19SB force field was applied to model the systems^[31]^. The initial structures were solvated in a truncated octahedral transferable intermolecular potential three point (termed as “TIP3P”) water box with a buffer of 10 Å around it. Then, counterions Na+ or Cl- were added to the systems for neutralization and 0.15 mol·L^-1^ NaCl was added to solvents.

After construction, the systems were firstly minimized for 15,000 cycles with a restraint of 500 kcal·mol^-1^·Å^-2^ on the RBD and ACE2. Then, all atoms encountered 30,000 cycles of minimization. Next, the systems were heated from 0 to 300 K in 300 ps and equilibrated for 700 ps with 10 kcal·mol^-1^·Å^-2^ positional restraint on non-solvent atoms. Finally, the WT, Omicron (Q493K) and Omicron (Q493R) simulation systems encountered 8 parallel rounds of 200 ns production MD simulations, respectively. During simulations, the temperature (300 K) and pressure (1 atm) was controlled by Langevin thermostat and Berendsen barostat, respectively. Long-range electrostatic interactions were treated by the Particle mesh Ewald algorithm with a grid size of 1 Å, and a cutoff of 10 Å was employed for short-range electrostatic and van der Waals interactions. The SHAKE algorithm was applied to restrain the bond with hydrogens. MD simulations were performed on Amber20.

### Markov State Model (MSM)

Starting with the code base of the current stable version 3.8.0 of MSMBuilder^[32]^, we developed a more robust algorithm to describe the transition process in Markov state model. Our algorithm modifies the fixed lag time into a random one by a kernel function, which is further used to count transition matrix and build MSM model. As a consequence, the MSM model based on our algorithm exhibits a more robust and powerful representation ability for describing the protein conformational space. We will discuss this method in-depth in our future publications.

From the trajectories of WT and Omicron (Q493R) system, the time-lagged independent component (tIC) analysis was firstly applied to decrease the dimension of the conformational space^[33]^. We selected the residue pairs, in which one residue was from RBD and the other was from ACE2. For each residue pair, we measured any pair of distances between the heavy atoms on RBD and those on ACE2 and kept the distances less than 5 Å as inputs for ContactFeaturizer. Then, the lag time of tIC analysis was set to 50 ns and 200 microstates were clustered by K-Centers algorithm. Multiple transition probability matrixes (TPMs) were further calculated according to the transitions among microstates. According to Eq. (1), the implied timescale test was performed to confirm the Markovian of microstates.

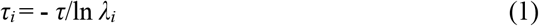

where *τ* represents the lag time for the TPMs, *λ_i_* is the i-th eigenvalue of the TPM and *τ*_i_ is the implied timescale for the i-th relaxation of the MSM. As a function of the lag time *τ, τ_i_* (especially τ1 for the slowest transition) is a constant when the transition among microstates are Markovian^[34,35]^. From the Markovian microstates, the macrostates were then clustered via the PCCA+ algorithm. Using transition path theory (TPT), the properties for transition, such as transition time and direction, were calculated^[36]^.

### MM/GBSA calculation

MMPBSA.py plugin in AmberTools20 was applied to exploit Molecular Mechanics/Generalized Born Surface Area (MM/GBSA) in binding free energy calculations in balanced trajectories for WT, Omicron (Q493K) and Omicron (Q493R) simulation systems^[37]^. In total, the binding free energy (ΔG_bind_) of RBD towards ACE2 is expressed as equation (2).

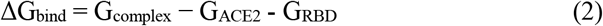

Meanwhile, the second law of thermodynamics reflects that ΔG_bind_ equals to enthalpy changes (ΔH) minus the product of entropy changes and temperature (TΔS), as equation (3) expresses.

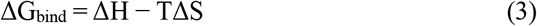

The system conformation entropy (termed as “-T△S”) is usually estimated by normal mode analysis with a quasi-harmonic model, however, accurate estimation of the conformation entropy for the protein-protein interactions remains challenging. Notably, the item could be omitted here considering that the differences of enthalpy (termed as “ΔH”) are large enough and the similarity of system conformation entropy among simulation systems^[38]^. Therefore, we omitted the calculation of the −TΔS term and only concentrated on the relative ordering of the free energy changes.

In the simulation process, ΔH is transformed into the sum of the molecular mechanical energies (ΔE_MM_) and the solvation free energy (ΔG_solv_), according to equation (4).

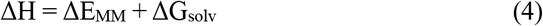

In addition, ΔE_MM_ consists of the intramolecular energy (ΔE_int_ including bond, angle and dihedral energies), van der Waals energy (ΔE_vdw_) and electrostatic energy (ΔE_ele_), while ΔG_solv_ consists of the polar (ΔG_P_) and the non-polar items (ΔG_np_). Equations (5) and (6) represent them.

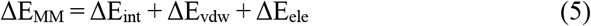

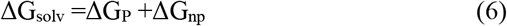

The Generalized Born model was used to calculate ΔG_P_. ΔG_np_ was calculated based on the solvent-accessible surface area (SASA) in equation (7).

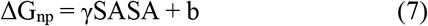

The solvation parameter γ and b were 0.00542 kcal (mol ^−1^·Å ^2^) and 0.92 kcal/mol, respectively.

The decomposition of the free energy into residues was subsequently carried out by the MMPBSA.py plugin^[37]^. During the decomposition process, 1-4 interactions were added to electric interactions and van der Walls interactions.

## Acknowledgments

We thank the SSRF BL02U1 and BL10U2 beam line for data collection and processing. We thank the X-ray crystallography platform of the Tsinghua University Technology Center for Protein Research for providing the facility support. This work was supported by funds from the National Natural Science Foundation of China (32171202), Tsinghua University Spring Breeze Fund (grant number 2020Z99CFY031), the National Science Foundation for Distinguished Young Scholars (32100973), the China Postdoctoral Science Foundation (grant number 2020M670295).

## Author contributions

X.W., T.W., J.G., and L.Z. conceived and designed the study. J.L. carried out protein purification, crystallization, diffraction data collection and structure determination with the help of H.Z., S.F. and Q.W.. Y.R. carried out SPR experiments. X.H. and T.W. conducted the MD simulation, Markov State Model construction, and binding free energy calculation. Y.Z. and D.L. synthesized gene of Omicron RBD. J.L., Y.R., Z.W. analyzed the structural data and made figures. J.G., T.W. and X.W. wrote the manuscript.

## Data availability statements

The coordinates and structure factors for the Omicron RBD-ACE2 complex were deposited in Protein Data Bank with accession code 7WHH.

