## extended table 1-3, extended figure 1-5 for "Structural and computational insights into the SARS-CoV-2 Omicron RBD-ACE2 interaction"

**Extended Data Table 1 | Data collection and refinement statistics**

| | Omicron RBD - ACE2 |
|---|---|
| **Data collection** | |
| Space group | $P4_12_12$ |
| Cell dimensions | |
| a, b, c (Å) | 104.71, 104.71, 227.10 |
| α, β, γ (°) | 90, 90, 90 |
| Resolution (Å) | 50-2.60(2.66-2.60) |
| $R_{merge}$ | 0.26 (2.45) |
| $I$/s$I$ | 11.3 (1.3) |
| Completeness (%) | 99.77(98.58) |
| Redundancy | 14.1 (11.2) |
| **Refinement** | |
| Resolution (Å) | 36.54-2.60 |
| No. reflections | 39525 |
| $R_{work}$/$R_{free}$ | 19.2/23.1 |
| No. atoms | |
| Protein | 6440 |
| Ligand/ion | 96 |
| Water | 131 |
| B-factors | |
| Protein | 50.47 |
| Ligand/ion | 90.06 |
| Water | 45.36 |
| R.m.s. deviations | |
| Bond lengths(Å) | 0.008 |
| Bond angles (°) | 0.93 |
| Ramachandran | |
| Favored (%) | 96.95 |
| Allowed (%) | 3.05 |
| Outliers (%) | 0.00 |

**Extended Data Table 2 | Contact residues of the WT and Omicron RBD-ACE2 interfaces**

| ACE2 | WT RBD | Omicron RBD |
|---|---|---|
| S19 | | A475, N477 |
| Q24 | A475, N487 | A475, N477, N487 |
| T27 | F456, A475, Y489 | F456, Y489 |
| F28 | Y489 | Y489 |
| D30 | K417, F456 | |
| K31 | Y489, Q493 | F456, Y489 |
| H34 | Y453, L455, Q493 | Y453, K493, S494 |
| E35 | Q493 | K493 |
| E37 | Y505 | |
| D38 | Y449 | Y449, S496, R498 |
| Y41 | Q498, T500, N501 | R498, T500, Y501 |
| Q42 | G446, Y449, Q498 | Y449, R498 |
| L79 | F486 | F486 |
| M82 | F486 | F486 |
| Y83 | F486, N487, Y489 | F486, N487, Y489 |
| N330 | T500 | T500 |
| K353 | G496, N501, G502, Y505 | Y501, G502, H505 |
| G354 | G502 | G502, H505 |
| D355 | T500 | T500 |
| R357 | T500 | T500 |
| R393 | Y505 | |

A distance cut-off of 4 Å was used.

**Extended Data Table 3 | The hydrogen bonds and salt bridges at the WT and Omicron RBD-ACE2 interfaces**

| | WT RBD | Length(Å) | ACE2 | Length(Å) | Omicron RBD |
|---|---|---|---|---|---|
| | | | S19(O) | 3.2 | N477(ND2) |
| | | | S19(OG) | 3.1 | A475(O) |
| | | | S19(N) | 3.4 | N477(OD1) |
| | N487(ND2) | 2.6 | Q24(OE1) | 2.8 | N487(ND2) |
| | K417(NZ) | 3.0 | D30(OD2) | | |
| | | | H34(ND1) | 2.9 | Y453(OH) |
| | Q493(NE2) | 2.8 | E35(OE2) | 3.1 | K493(NZ) |
| | Y505(OH) | 3.2 | E37(OE2) | | |
| | | | D38(OD1) | 2.9 | R498(NH1) |
| **Hydrogen bonds** | | | D38(OD1) | 2.8 | S496(OG) |
| | Y449(OH) | 2.7 | D38(OD2) | 2.5 | Y449(OH) |
| | T500(OG1) | 2.6 | Y41(OH) | 2.6 | T500(OG1) |
| | N501(N) | 3.7 | Y41(OH) | | |
| | G446(O) | 3.3 | Q42(NE2) | | |
| | Y449(OH) | 3.0 | Q42(NE2) | 3.4 | Y449(OH) |
| | Y489(OH) | 3.5 | Y83(OH) | 3.5 | Y489(OH) |
| | N487(OD1) | 2.7 | Y83(OH) | 2.4 | N487(OD1) |
| | G502(N) | 2.8 | K353(O) | 2.7 | G502(N) |
| | Y505(OH) | 3.7 | R393(NH2) | | |
| | K417(NZ) | 3.9 | D30(OD1) | | |
| | K417(NZ) | 3.0 | D30(OD2) | | |
| **Salt bridges** | | | E35(OE2) | 3.1 | K493(NZ) |
| | | | D38(OD1) | 2.9 | R498(NH1) |
| | | | D38(OD1) | 3.7 | R498(NH2) |

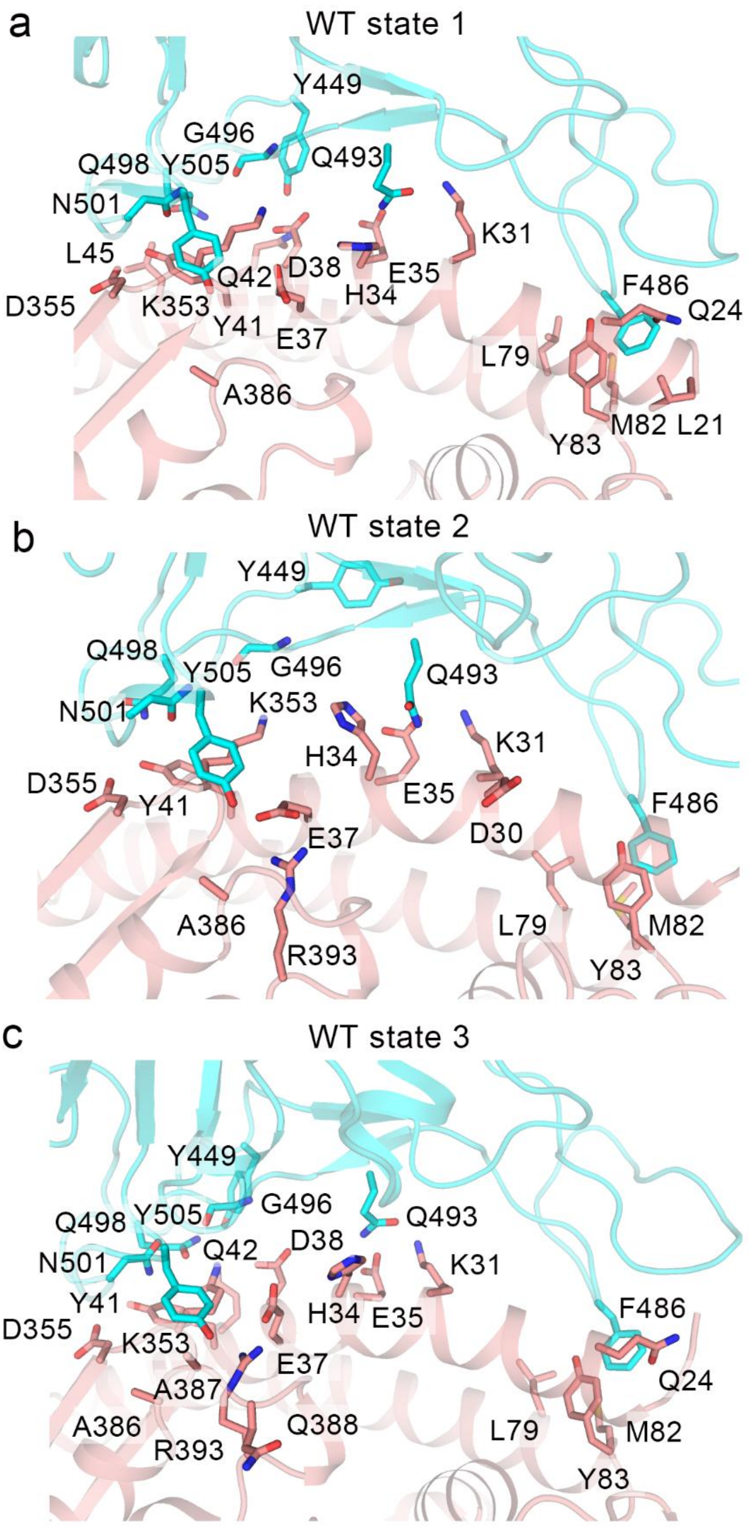

**Extended Data Fig. 1** The interactions between ACE2 and RBD in different WT states. The residues in ACE2 within 5 Å to Y449, F486, Q493, G496, Q498, N501, and Y505 are shown in sticks. The RBD of WT and ACE2 are shown in cyan and salmon, respectively.

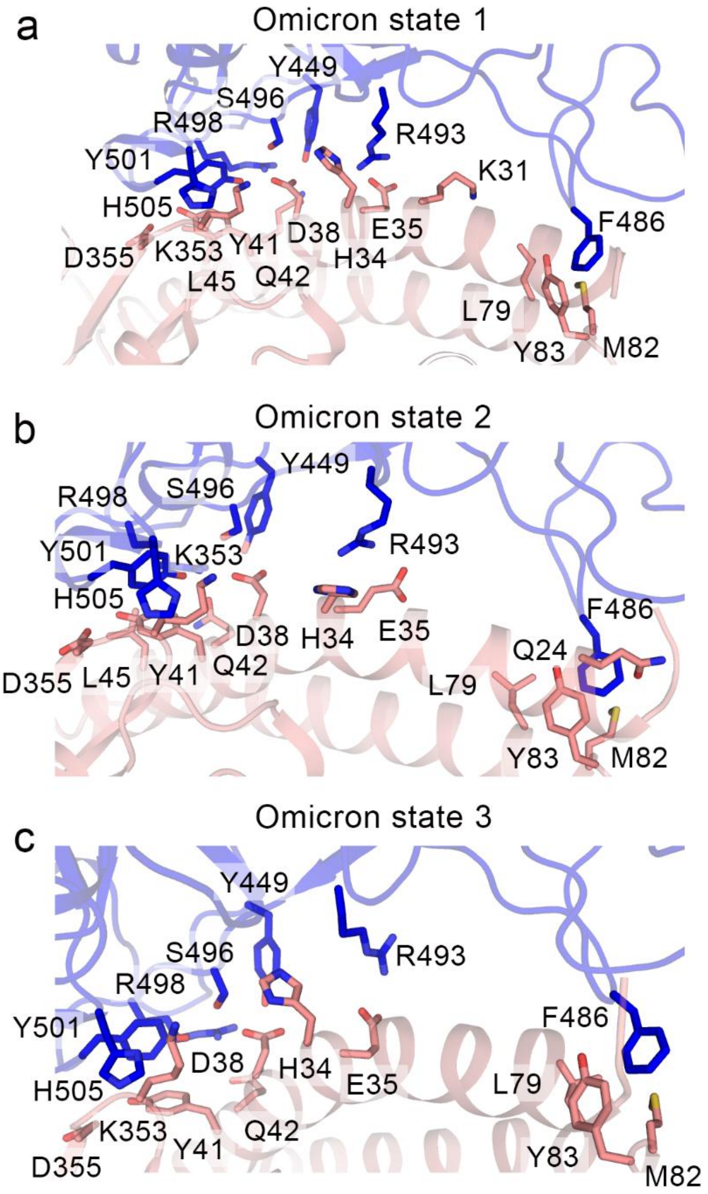

**Extended Data Fig. 2** The interactions between ACE2 and RBD in different Omicron states. The residues in ACE2 within 5 Å to Y449, F486, R493, S496, R498, Y501, and H505 are shown in sticks. The RBD of Omicron and ACE2 are shown in blue and salmon, respectively.

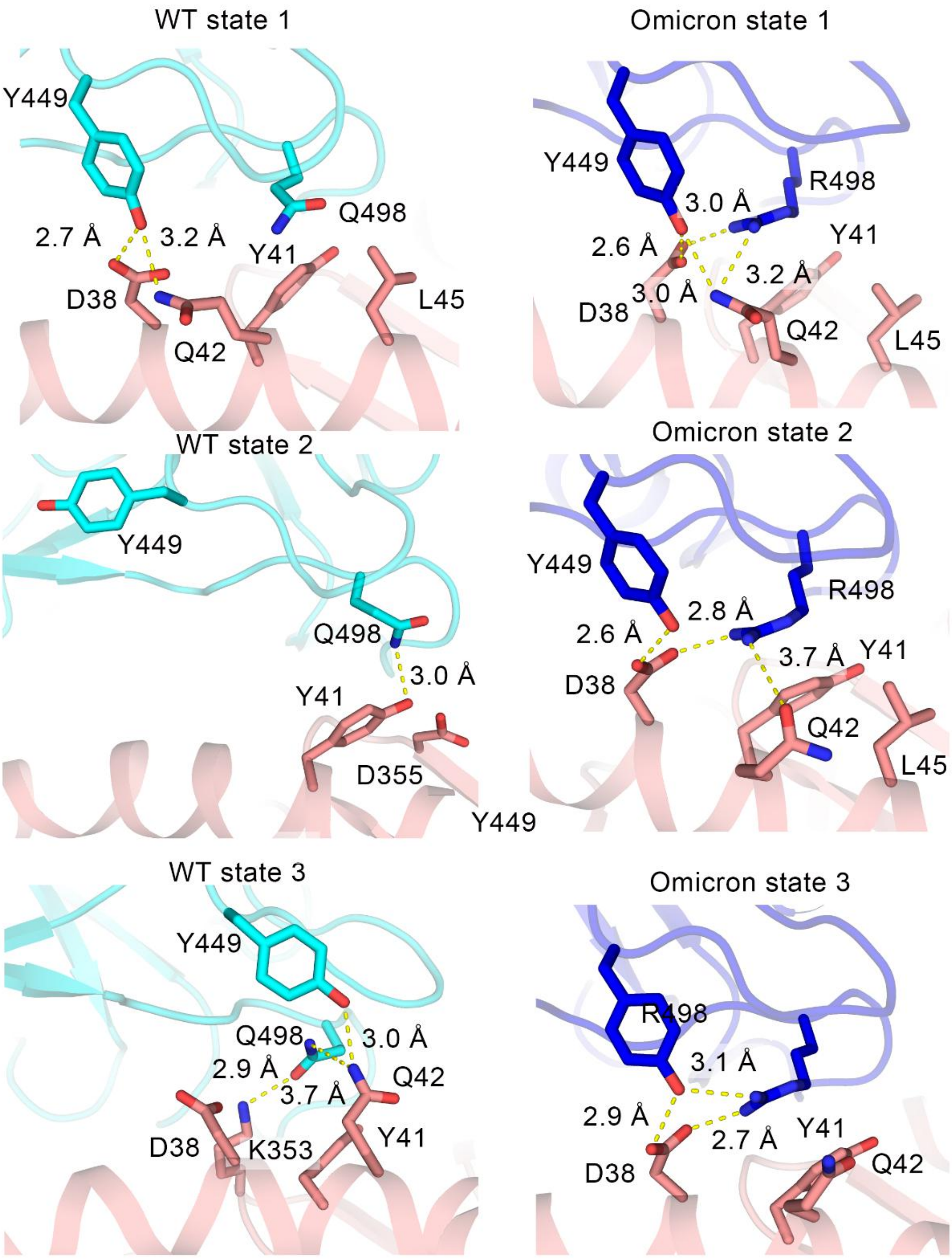

**Extended Data Fig. 3** The interactions between Y449 and Q/R498 of RBD with ACE2 in WT and Omicron systems. The residues in ACE2 within 5 Å to Y449 and Q/R498 are shown in sticks while interactions are shown in yellow dashed lines. The RBD of WT, the RBD of Omicron and ACE2 are shown in cyan, blue and salmon, respectively. In WT system, Y449 shows a unique horizontal pose in state 2 which is unseen in other states. The loss of interactions with D38 and Q42 may lead to the decrease of Y449 contribution in WT state 2. In the Omicron system, however, the hydrogen bond network among D38, Q42 Y449 and R498 reduces the possibility of the horizontal conformation of Y449 and also promotes the interactions between R498 and ACE2. In WT state 3, Q498 interacts with Q42, which similarly promotes its binding affinity ($\Delta G$ = -5.43 ± 2.35 kcal/mol).

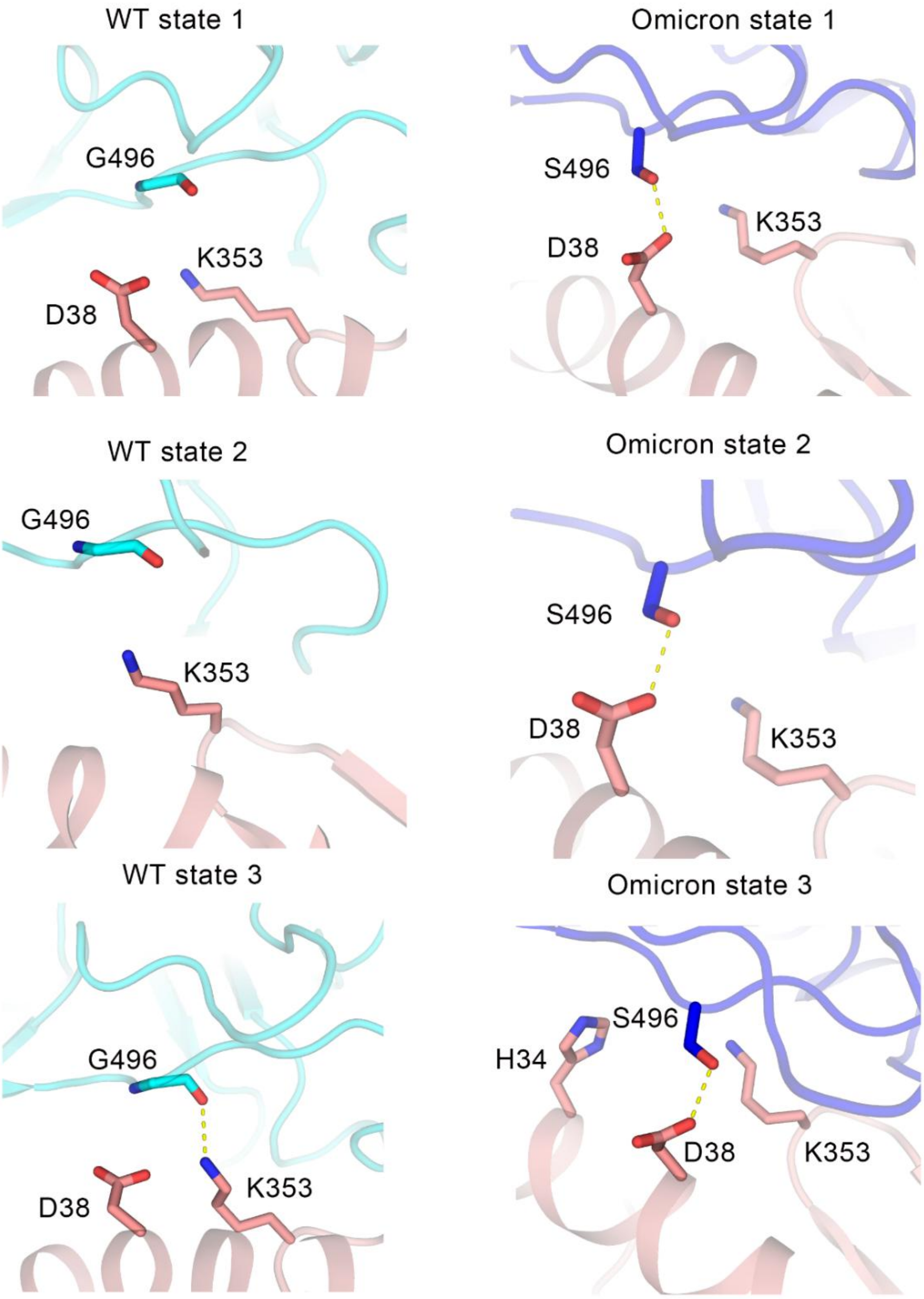

**Extended Data Fig. 4** The interactions between G/S496 of RBD with ACE2 in WT and Omicron systems. The residues in ACE2 within 5 Å to G/S496 are shown in sticks while interactions are shown in yellow dashed lines. The RBD of WT, the RBD of Omicron and ACE2 are shown in cyan, blue and salmon, respectively. In WT system, G496 in state 1 and state 2 show no interaction with ACE2, leading to weak contribution to the binding affinity (-0.95 ± 1.18 and -0.01 ± 0.11 kcal/mol, respectively). As for state 3, G496 interacts with K353 in its main chain, which leads to an increase of the binding ability (-2.05 ± 1.11 kcal/mol). As a contrast, in Omicron system, S496 has interaction with D38 in all three macrostates and behaves a lower binding free energy.

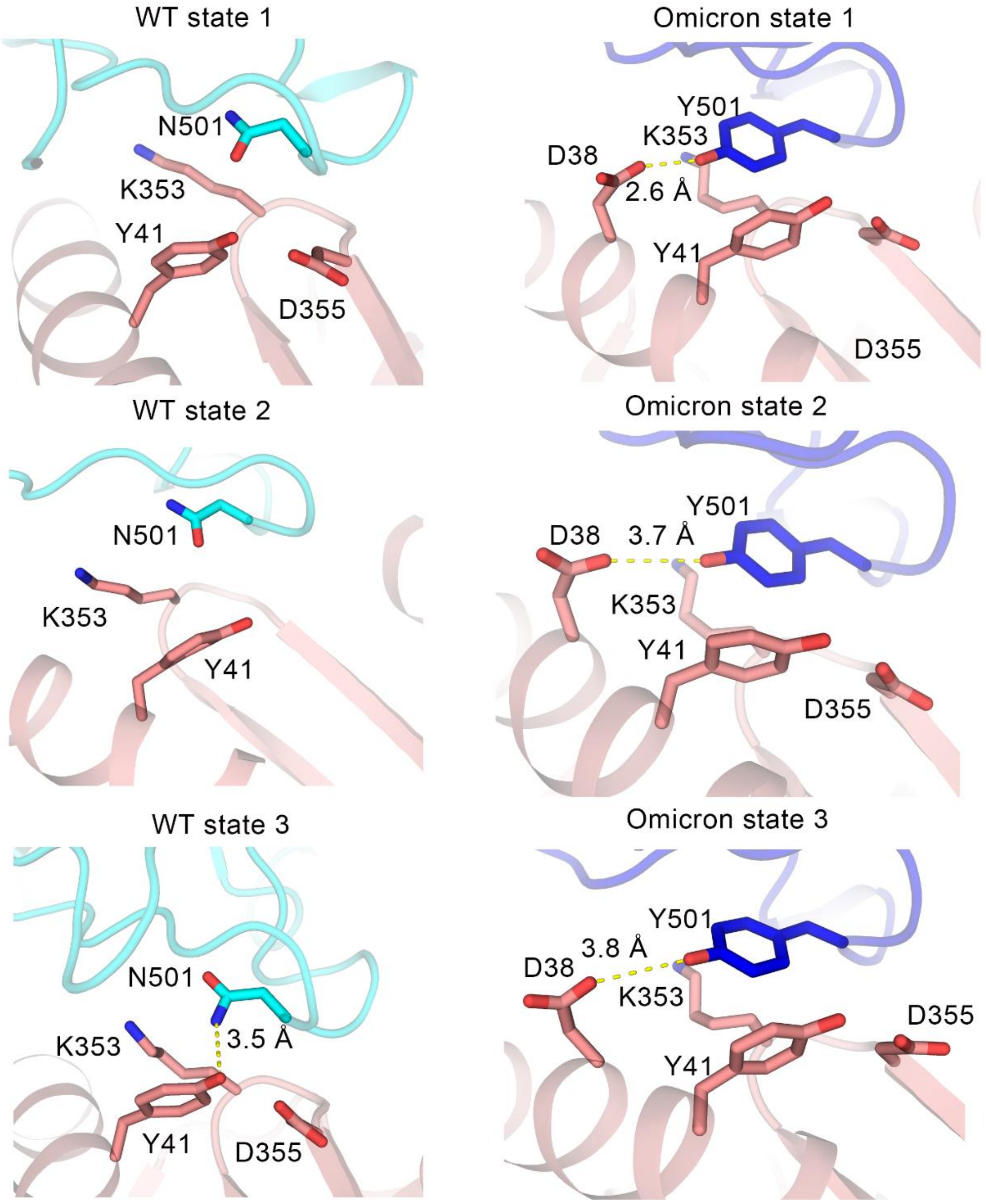

**Extended Data Fig. 5** The interactions between N/Y501 of RBD with ACE2 in WT and Omicron systems. The residues in ACE2 within 5 Å to N/Y501 are shown in sticks while interactions are shown in yellow dashed lines. The RBD of WT, the RBD of Omicron and ACE2 are shown in cyan, blue and salmon, respectively. In WT system, N501 also has no interaction with ACE2 in states 1 and 2 and shows weak energy contribution (-1.14 ± 0.79 and -0.74 ± 0.77 kcal/mol, respectively). Even in WT state 3, the energy does not decrease too much with the hydrogen bond to Y41 (-2.61 ± 1.28 kcal/mol). This phenomenon may be caused by the hydrophilic sidechain of N501 embedded in a hydrophobic environment made by the sidechains of K353 and Y51. As N501Y changes to a residue with a longer sidechain and a hydrophobic phenyl ring in Omicron system, the interaction with D38 is stable in each macrostate and the ring of tyrosine is suitable in the hydrophobic environment.
